# Long-read Individual-molecule Sequencing Reveals CRISPR-induced Genetic Heterogeneity in Human ESCs

**DOI:** 10.1101/2020.02.10.942151

**Authors:** Chongwei Bi, Lin Wang, Baolei Yuan, Xuan Zhou, Yu Pang, Li, Sheng Wang, Yuhong Xin Gao, Yanyi Huang, Mo Li

## Abstract

Accurately quantifying the genetic heterogeneity of a cell population is essential to understanding of biological systems. We develop a universal method to label individual DNA molecules for analyzing diverse types of rare genetic variants, with frequency as low as 4×10^−5^, using short- or long-read sequencing. It enables base-resolution haplotype-resolved quantitative characterization of rare variants. It provides the first quantitative evidence of persistent nonrandom large deletions and insertions following DNA repair of double-strand breaks induced by CRISPR-Cas9 in human pluripotent stem cells.

---

Molecular consensus sequencing has been developed to enhance the accuracy of next-generation sequencing (NGS) using unique molecular identifier (UMI)(*1-3*). The main concepts of this strategy include barcoding each molecule before amplification, and correcting sequencing error using the consensus sequence of reads containing the same barcode, and hence eliminating the random errors introduced by sequencing chemistry or detection. However, current strategies are inadequate for many types of sequences especially the large structural variants or repetitive sequences (Fig. S1). Single molecule sequencing technologies can better resolve complex genetic variants by providing long reads, but they have a lower raw read accuracy(*3*). To overcome these limitations, we have developed a strategy termed targeted <u>I</u>ndividual <u>D</u>NA <u>M</u>olecule <u>seq</u>uencing (IDMseq). IDMseq guarantees that each original DNA molecule is uniquely represented by one UMI group (a set of reads sharing the same UMI) after sequencing, thus preventing false UMI groups and allowing quantification of allele frequency in the original population (Fig. S1 & 2a). It is designed to be adaptable to various sequencing platforms, and combines error correction by molecular consensus with long-read sequencing, thus enabling sensitive detection of all classes of genetic variants, including SNVs, indels, large deletions, and complex rearrangements.

To verify that IDMseq can detect subclonal variants below the sensitivity limit of NGS (∼1%(*4, 5*)), we constructed synthetic cell populations harboring a mutation at various pre-determined allele frequencies. We knocked in a homozygous single nucleotide variant (SNV) in the *EPOR* gene using CRISPR-Cas9 in the H1 human embryonic stem cells (hESCs)(Fig. S3a-c). A rare subclonal mutation in a population of cells is simulated by admixing the genome of knock-in and wild-type cells at different ratios.

First, we tested if IDMseq could overcome the high base-calling error of Nanopore sequencing in rare mutation detection. A 168 bp stretch of DNA encompassing the knock-in SNV was labeled with UMIs and amplified from a population with the ratio of 1:100 between knock-in and wild-type alleles. We developed a bioinformatic toolkit called <u>V</u>ariant <u>A</u>nalysis with <u>U</u>MI for <u>L</u>ong-read <u>T</u>echnology (VAULT) to analyze the sequencing data (Fig. S2b, see methods). The results showed that 36.5% of reads contained high-confidence UMI sequences (Table 1). Based on a pre-set threshold of a minimum of 5 reads per UMI group, those reads are binned into 284 UMI groups. It is worth noting that every UMI group represents an original allele in the genome of the initial population. VAULT analysis showed that 2 UMI groups contained the knock-in SNV (Fig. S4b). Furthermore, no spurious mutation was detected. Importantly, when the trimmed reads were pooled for variant analysis without considering UMIs, no variant could be detected by the same algorithms, proving the superior sensitivity afforded by IDMseq. These results suggest that IDMseq on the single-molecule Nanopore sequencing platform is able to accurately call rare variants without false positives.

**Table 1.** Summary of individual sequencing runs

| Gene | Mutant allele frequency (%) | Amplicon size | Sequencing platform | Read count | Reads with UMI | UMI groups for variant calling (>= 5 reads) | UMI groups with introduced mutation | Somatic SNV count | Somatic SNV load Per megabase | SV groups |
| --- | --- | --- | --- | --- | --- | --- | --- | --- | --- | --- |
| *EPOR* | 1:100 (1%) | 168 bp | Nanopore | 17,634 | 6,444 | 284 | 2 (0.7%) | 0 | N/A | N/A |
| *EPOR* | 1:1,000 (0.1%) | 6,789 bp | PacBio | 227,206 | 136,399 | 3,184 | 4 (0.126%) | 273 | 8.9 | 3 |
| *EPOR* | 1:10,000 (0.01%) | 168 bp | Nanopore | 1,093,683 | 494,009 | 15,598 | 1 (0.006%) | 10 | 3.8 | N/A |
| *EPOR* | 1:10,000 (0.01%) | 168 bp | Illumina | 7,488,257 | 7,236,007 | 132,341 | 5 (0.004%) | 85 | 3.9 | N/A |
| *PANX1* | N/A | 7077 bp | Nanopore | 2,761,805 | 613,147 | 3,566 | N/A | 293 | 11.6 | 200 (5.6 %) |
| *PANX1* | N/A | 6595 bp | Nanopore | 3,078,165 | 1,042,582 | 8,870 | N/A | 843 | 14.4 | 232 (2.6 %) |

Detection of rare variants in clinical settings often demands sensitivities well below that of prevailing NGS platforms (ca. 10^−2^). For instance, early cancer detection using circulating tumor DNA is estimated to require a sensitivity of at least 1 in 10,000(*6*). To simulate this scenario, we next sequenced the same 168 bp region in a population with the ratio of 1:10,000 between knock-in and wild-type alleles. It is worth noting that the UMI-labeling reaction contained only around 5 copies of the knock-in allele. A 48-hour sequencing run on the MinION acquired 1.1 million reads (Fig. S4a). VAULT showed that 45.2% of reads contained high-confidence UMI sequences (Table 1). These reads were binned into 15,598 UMI groups of which one (0.6×10^−4^) contained the knock-in SNV (Fig. 1b). Ten other SNVs were also identified in ten UMI groups. We considered if these were PCR artifacts, as the main source of errors in UMI consensus sequencing originates from polymerase replication error in the barcoding step(*7*). The Platinum SuperFi DNA polymerase we used has the highest reported fidelity (>300X that of *Taq* polymerase). It not only significantly reduces errors in the barcoding and amplification steps, but also captures twice more UMIs in the library than *Taq*(*7*). Theoretically, Platinum SuperFi polymerase introduces ∼6 errors in 10^6^ unique 168-bp molecules in the UMI-labeling step. Accordingly, this type of inescapable error is expected to be around 0.09 in 15,598 UMI groups, and thus cannot account for the observed SNV events. This lets us to conclude that the ten SNVs are rare somatic mutations that reflect the genetic heterogeneity of hESCs as described previously(*8*). These data provided an estimate of 3.8 somatic SNVs per megabase (Mb), which is consistent with the reported frequency of somatic mutation in coding sequence in normal healthy tissues(*9*).

**Fig. 1.**
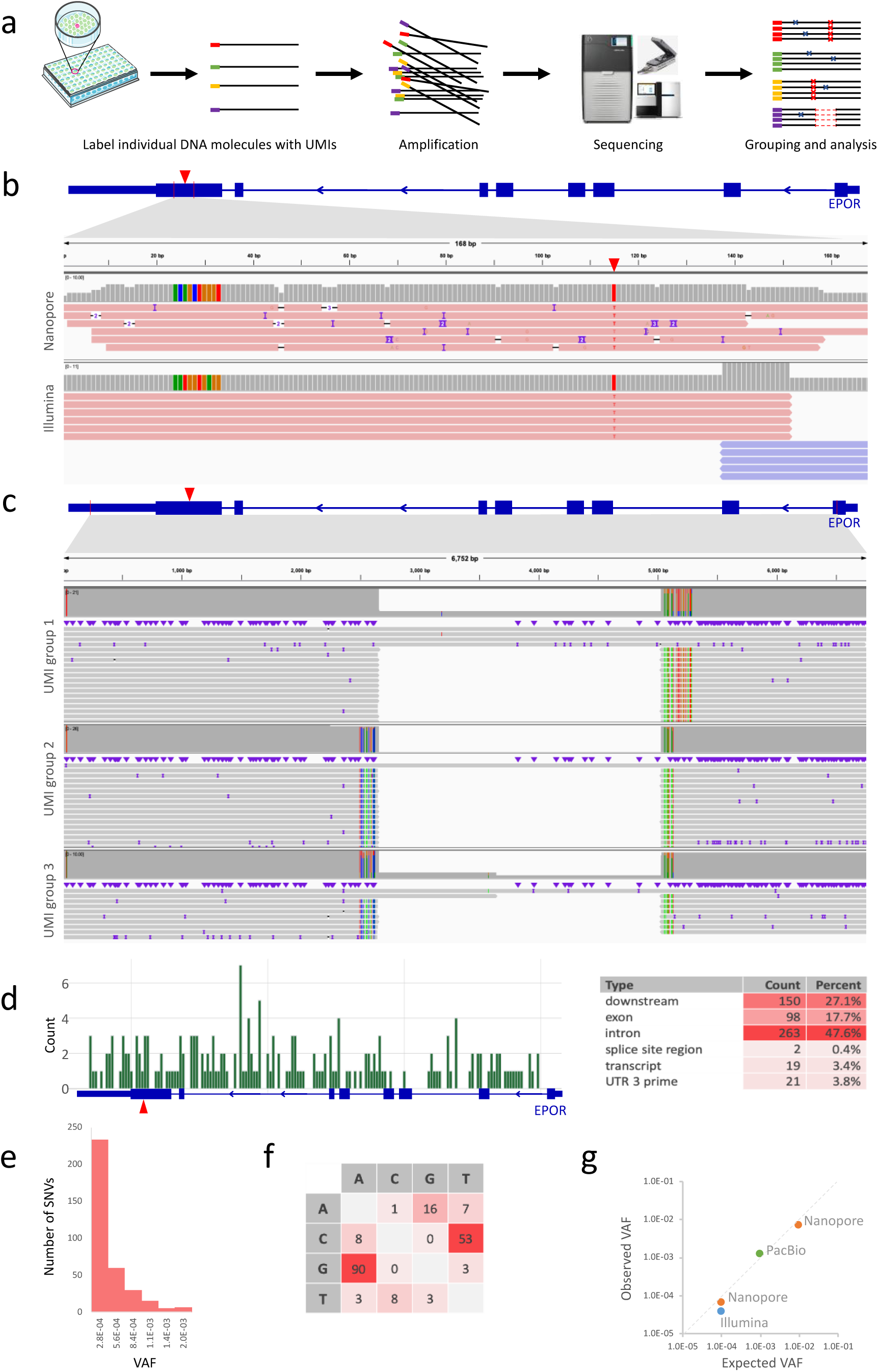
IDMseq for detection of subclonal variants. **a**, Schematic representation of IDMseq. Individual DNA molecules are labeled with unique UMIs and amplified for sequencing on appropriate platforms (e.g. Illumina, PacBio, and Nanopore). During data analysis, reads are binned by UMIs to correct errors introduced during amplification and sequencing. Both SNV and SV calling are included in the analysis pipeline. **b**, Examples of Integrative Genomics Viewer (IGV) tracks of UMI groups in which the spike-in SNV in the 1:10000 population was identified by IDMseq and VAULT. The knock-in SNV is indicated by the red triangle in the diagram of the EPOR gene on top, and also shown as red “T” base in the alignment map. The gray bars show read coverage. The ten colored bars on the left side of the coverage plot represent the UMI sequence for the UMI group. Individual Nanopore (top) and Illumina (bottom) reads within the group are shown under the coverage plot. **c**, Large SVs detected by IDMseq in the 1:1000 population on the PacBio platform. Three UMI groups are shown with the same 2375bp deletion. Group 1 represents one haplotype, and Group 2&3 represent a different haplotype. Colored lines represent the SNPs detected in each group. Thick blue boxes: exons; think blue boxes UTRs. Thin vertical red lines in the gene diagram represent PCR primer location. **d**, Distribution of SNVs detected by PacBio sequencing in conjunction with IDMseq and VAULT. One of the SNVs was also found in the Nanopore dataset. The spike-in SNV (1:1000) is indicated by the red triangle. The table on the right summarizes the frequency of SNVs in different annotation categories. **e**, Frequency distribution of the variant allele fraction of SNVs detected by IDMseq in PacBio sequencing of the EPOR locus. **f**, The spectrum of base changes among somatic SNVs. The majority of base changes are G to A and C to T. **g**, Comparison between observed VAF and expected VAF in different experiments and sequencing platforms.

The length of 168-bp amplicon also allowed benchmarking against the industry standard Illumina sequencing, which features shorter reads but higher raw-read accuracy. We then sequenced the same 1:10,000 mixed population on an Illumina MiniSeq sequencer and obtained 7.5 million paired-end reads (Fig. S4a). The results showed that 96.6% of reads contained high-confidence UMI sequences that were binned into 132,341 UMI groups, in which 5 (4×10^−5^) contained the knock-in SNV (Table 1, Fig. 1b). The calculated somatic SNV load was 3.9 per Mb, which closely matches the Nanopore data.

We next applied IDMseq to a larger region (6,789 bp) encompassing the knock-in SNV in a population with 0.1% mutant cells on a PacBio platform (Fig. S4a). VAULT showed that 60.0% of the high-fidelity long reads contain high-confidence UMIs, binned into 3,184 groups. Four UMI groups (1.26×10^−3^) contained only the knock-in SNV. Another 186 groups contained 273 SNVs (174 groups with 1 SNV, 9 groups with 2 SNVs, and 3 groups with 27 SNVs, Table 1). Again, polymerase error during barcoding (∼0.82 error in 3,184 UMI groups) cannot account for the observed SNVs, suggesting that most SNVs are true variants. Interestingly, structural variant (SV) analysis showed that the three groups with 27 SNVs shared the same 2,375 bp deletion. Haplotyping using the SNVs revealed that the three groups came from two haplotypes (Fig. 1c). This large deletion is far away from the Cas9 target site and thus less likely the result of genome editing. After excluding the SNVs in the large-deletion alleles, the remaining 192 SNVs distributed evenly in the region (Fig. 1d). Functional annotation of the SNVs showed that 17 of 192 caused an amino acid change. The spectrum of base changes and distribution of variant allele frequency (VAF) are consistent with published work(*9*) (Fig. 1e, f). These data provide an estimate of about 8.9 somatic SNVs per Mb.

Taken together these data showed that IDMseq provides reliable detection of rare variants (at least down to 10^−4^) and accurate estimate of variant frequency (Fig. 1g). It is useful for characterizing the spectrum of somatic mutations in human pluripotent stem cells (hPSCs). Furthermore, it revealed a previously unappreciated phenomenon of spontaneous large deletion in hPSCs. Due to its large size and low frequency (VAF≈0.1%), this SV would have been missed by short-read sequencing or ensemble long-read sequencing. Yet, it is conceivable that such an SV could confer growth advantage to the cell carrying it, and therefore has implications for the safety of hPSC in clinical settings. These findings nicely demonstrate the power of the combination of long-read sequencing and IDMseq in resolving complex genetic heterogeneity.

Despite its widespread adoption of the CRISPR-Cas9 system as an efficient and versatile genome-editing tool, the impact of CRIPSR on human genome integrity remains poorly understood(*10-12*). Previous work indicated that the most prevalent DNA repair outcomes after Cas9 cutting are small indels (typically < 20 bp)(*13, 14*). Unexpectedly, recent studies revealed large and complex SVs over several kilobases represent a significant portion of the on-target mutagenesis effect of Cas9(*10, 15*). Importantly, to date, the analysis of large-deletion alleles came either from ensemble amplicon sequencing(*10, 15*) or whole-genome sequencing(*15*). The former is prone to amplification bias, and the latter cannot adequately detect large and complex variants due to the limited read length. Thus, we applied IDMseq to hESCs following CRISPR-Cas9 editing, to offer an unbiased quantification of the frequency and molecular feature of the DNA repair outcomes of double-strand breaks induced by Cas9.

We targeted exon 1 (Pan1) and exon 3 (Pan3) of the Pannexin 1 (*PANX1*) gene with two efficient gRNAs (Fig. 2a, Fig. S5e). Forty-eight hours after electroporation of Cas9 complexed with the Pan1 or Pan3 gRNA, H1 ESCs were harvested for IDMseq. The surveyed region is 7,077 bp for Pan1 and 6,595 bp for Pan3. A 48h Nanopore sequencing run yielded 2.8 million and 3.1 million reads for Pan1 and Pan3, which were binned into 3,566 and 8,870 UMI groups, respectively (Table 1, Fig. 2b, Fig. S5a).

**Fig. 2.**
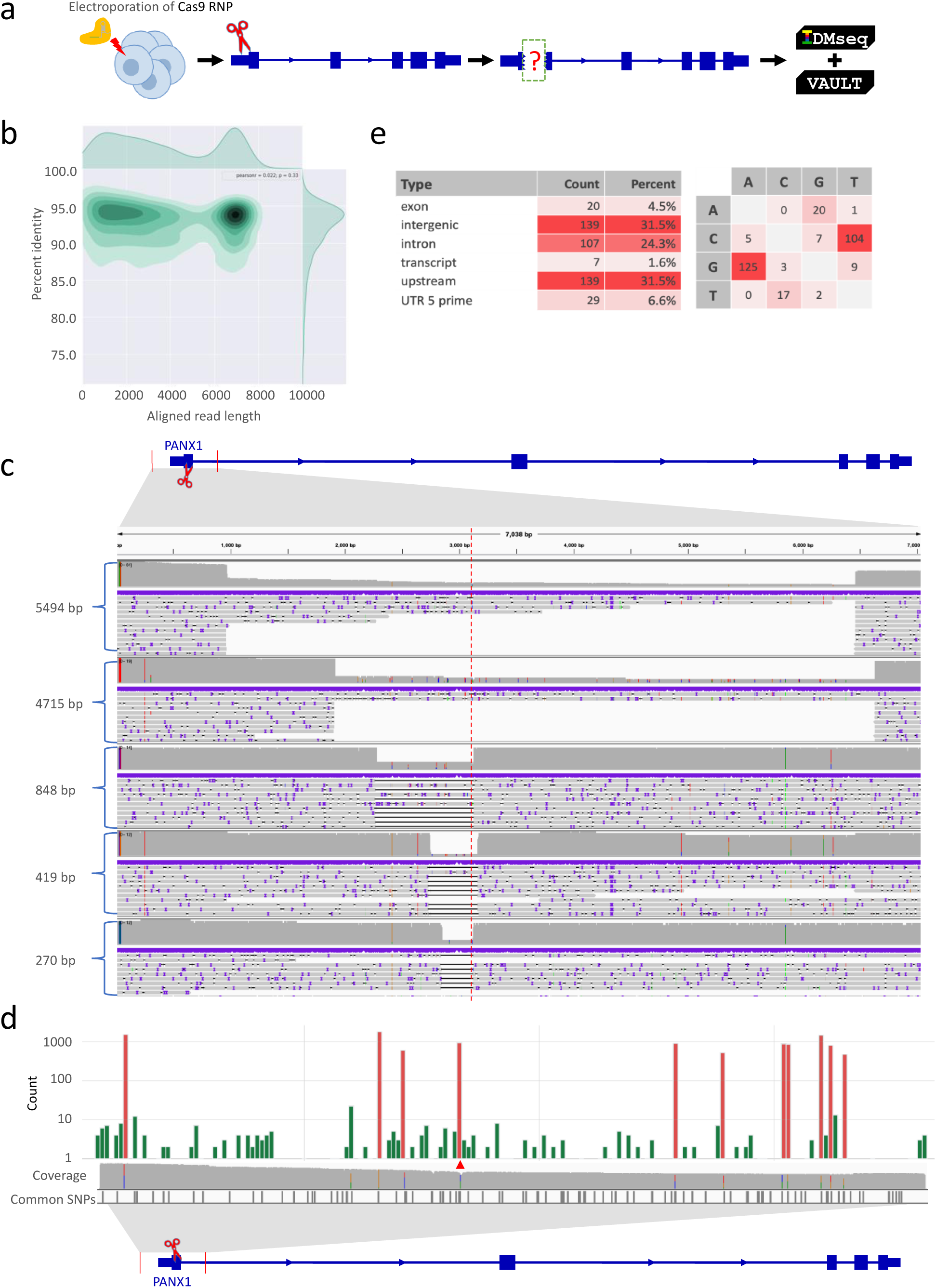
Quantitative analysis of DNA repair outcome of Cas9-induced DNA double-strand break in hESCs. **a**, Schematic representation of the experimental design. Cas9 RNPs designed to cleave the first exon of *PANX1* were electroporated to H1 hESCs. IDMseq was used to analyze the locus in edited hESCs 48 hours later. **b**, Aligned read length vs. percent identity plot using kernel density estimation of Nanopore sequencing data of a 7077 bp region encompassing the Cas9 cleavage. **c**, Large SVs detected by IDMseq and VAULT in edited hESCs. Five SV groups were shown with deletion length ranging from 270 bp to 5494 bp. The red dotted line represents the Cas9 cutting site. The coverage of Nanopore reads is shown on top of each track in gray. The colored lines on the left side of the coverage plot represent the UMI for the group. Individual Nanopore reads within the group are shown under the coverage plot. **d**, Distribution of SNVs detected by IDMseq and VAULT in edited hESCs. Somatic SNVs are shown in green, while the cell-line specific SNVs are shown in red. Somatic SNVs cannot be detected if variant calling is done en masse without UMI analysis (see the coverage track). Cell-line specific SNVs are detected in ensemble analysis (see colored lines in the coverage track) and most of them have been reported as common SNPs in dbSNP-141 database (Common SNPs track). The Cas9 cut site is indicated by the red triangle. **e**, Analysis of somatic mutations detected in CRISPR-edited hESCs based on functional annotation and base change. The majority of base changes are G to A and C to T.

We first surveyed SVs (>30 bp) in UMI groups. After SV calling and filtering out lowly supported SVs (see methods), 200 (5.6 %) of the 3,566 UMI groups contained 200 SVs in Pan1-edited cells, including 195 deletions and 5 insertions. The size of SVs ranged from 31 to 5,506 bp (Fig. 2c, Fig. S6a). Intriguingly, some large deletions were independently captured multiple times. For example, 56 (28.0%) UMI groups have the same 5,494-bp deletion and 18 (9.0%) UMI groups have the same 4,715-bp deletion. For the insertion variants, 3 of the 5 UMI groups shared the same SV.

When a different gRNA (Pan3) was used, 232 (2.6%) of 8,870 UMI groups contained 240 SVs (178 deletions, 50 insertions and 12 inversions), with size ranging from 31 to 4,238 bp (Fig. S6b). Importantly, reoccurring SVs were also detected with Pan3. Twenty-seven (32.1%) UMI groups shared the same 4,238-bp deletion, and 6 (2.5%) groups shared a 2,750-bp insertion. These data provided the first quantitative evidence that the repair outcome of Cas9 cutting is not random and there are likely hotspots for Cas9-induced large deletions or insertions.

We next analyzed SNVs in these two data sets. Cas9 editing with the Pan1- and Pan3 gRNAs resulted in similar SNV patterns (Fig. 2d, Fig. S5f). The results of Pan1 showed that 2,731 (76.6%) of 3,566 UMI groups contained 10,871 SNPs, while for Pan3 8,018 (90%) of 8,870 UMI groups contained 23,477 SNVs. In both cases, the SNVs fell into two frequency ranges. Most SNVs in the high-frequency category (red in Fig. 2d & Fig. S5f) have been reported in the common SNP database. The low-frequency SNVs (green in Fig. 2d & Fig. S5f) distributed evenly in the locus and did not overlap with known SNPs, likely representing somatic mutations. There was no obvious enrichment of SNVs around the cutting site, which is consistent with previous reports(*16*). The frequency of presumed somatic mutations for Pan1 (293 somatic mutations) and Pan3 (843 somatic mutations) is 11.6 and 14.4 per Mb, respectively, which is slightly higher than the average value (∼7/Mb) reported by NGS(*17*). The spectrum (Fig. 2d-e, Fig. S5f) and VAF (Fig. S6c-e) of single nucleotide substitutions were consistent with published data(*17*).

Besides SNVs and SVs, VAULT also reported many small indels around the Cas9 cleavage site. We compared the indels with the Sanger sequencing data of single-cell derived clones. The results showed that IDMseq correctly identified a subset of the deletion alleles (Fig. S5b-e).

In summary, IDMseq and VAULT enable quantitation and haplotyping of both small and large genetic variants at the subclonal level. They are easy to implement and compatible with all current sequencing platforms, including the portable Oxford Nanopore MinION. IDMseq provides an unbiased base-resolution characterization of on-target mutagenesis induced by CRISPR-Cas9, which could facilitate the safe use of the CRISPR technology in the clinic. The high sensitivity afforded by IDMseq and VAULT may be useful for early cancer detection using circulating tumor DNA or detection of minimal residual diseases. Our results showed that IDMseq is accurate in profiling rare somatic mutations, which could aid the study of genetic heterogeneity in tumors or aging tissues. IDMseq in its current form only sequences one strand of the DNA duplex, and its performance may be further improved by sequencing both strands of the duplex.

## Materials and Methods

### Generation of the knock-in hESC line

The H1 hESC line was purchased from WiCell and cultured in Essential 8™ medium (ThermoFisher) on hLaminin521 (ThermoFisher) coated plate in a humidified incubator set at 37°C and 5% CO2. Electroporation of CAS9 RNP was done using a Neon Transfection System (ThermoFisher) using the following setting: 1600 v/10 ms /3 pulses for 200,000 cells in Buffer R (Neon Transfection kit) premixed with 50 pmol Cas9 protein (CAT#M0646T, New England Biolabs), 50 pmol single guide RNA (sgRNA) and 30 pmol single-stranded oligodeoxynucleotides (ssODN, purchased from Integrated DNA Technologies, Inc.) template. After 48 hours, single cells were collected and seeded at 1,000 single cells per well (6-well format). Seven days later, single colonies were picked for passaging and genotyping.

### CRISPR-Cas9 editing of hESCs

CRISPR-Cas9 editing of the PANX1 locus in H1 hESCs were performed in the same way as the generation of knock-in hESCs except for the omission of the ssODN template. After 48 hours, cells are collected for the genome extraction and library preparation. The Pan1 sgRNA sequence is 5’ATCCGAGAACACGTACTCCG-TGG(PAM)3’ and Pan3 sgRNA is 5’GCTGCGAAACGCCAGAACAG-CGG(PAM)3’.

### UMI primer design

The UMI primer contains a 3’ gene-specific sequence, a UMI sequence, and a 5’ universal primer sequence. The 3’ gene-specific sequence is designed with the same principle as PCR primers. We chose the sequence with an annealing temperature higher than 65 °C to improve specificity to the target gene. The internal UMI sequence consists of multiple random bases (denoted by Ns). The number of random bases is determined by the number of targeted molecules. We chose a short UMI sequence (10-12 nt) to reduce the sequencing errors within the UMI. We adopted a unique sequence structure in the UMI (e.g. NNNNTGNNNN) to avoid homopolymers that may introduce errors due to polymerase slippage or low accuracy of Nanopore sequencing in these sequences. Several studies have also pointed out that both Illumina and PacBio are prone to errors in such regions18,19. The structured UMI design also serves as a quality control in the UMI analysis. The 5’ universal primer sequence is used to uniformly amplify all UMI tagged DNA molecules. It is designed to avoid non-specific priming in the target genome.

### UMI labeling

Genomic DNA is extracted using the Qiagen DNeasy Blood & Tissue Kit. The concentration is determined using a Qubit 4 Fluorometer (ThermoFisher). The UMI labeling step is done by one round of primer extension with a high-fidelity DNA polymerase. After UMI labeling DNA is purified by AMPure XP beads, followed by PCR amplification using the universal primer and the gene-specific reverse primer. This amplification will generate enough UMI-labeled DNA for downstream sequencing. In addition to one-ended labeling, two-ended UMI labeling can also be achieved by performing an additional UMI-labeling step with a reverse primer tagged with a UMI (Supplementary Fig. 2a). Two-ended UMI labeling could increase analyzable reads and provides extra benefit in accuracy. However, we found that due to the fact that UMI labeling is limited by primer efficiency, one-ended labeling will cover more molecules. Additional UMI-labeling and purification steps result in higher loss of DNA of interest. Since the procedure of one-ended labeling is simple and efficient, we used one-end UMI labeling for all experiments in this study.

### Library preparation and sequencing

For Nanopore sequencing, library preparations were done using the ligation sequencing kit (Cat# SQK-LSK109, Oxford Nanopiore Technologies). The sequencing runs were performed on an Oxford Nanopore MinION sequencer using R9.4.1 flow cells. Base calling of Nanopore reads was done using the official tool termed Guppy (v3.2.1). For PacBio sequencing, library preparations were done using the Sequel Sequencing Kit 3.0. The sequencing runs were performed by the BIOPIC core facility at Peking university (Beijing, China) on a PacBio Sequel using Sequel SMRT Cell 1M v3. HiFi Reads were generated by the official tool termed ccs (v3.4.1). All procedures were preformed according to manufacturer’s protocols. For Illumina sequencing, library preparations were performed using the NEBNext Ultra II DNA Library Prep Kit for Illumina. An unrelated RNA library prepared using the same kit was pooled to increase the complexsity of final library. The sequencing of paired-end 150bp reads was done on an Illumina Miniseq.

### Data processing

VAULT uses several published algorithms for UMI extraction, alignment, and variant calling. The whole analysis can be done with one command. In brief, Nanopore reads are trimmed to remove adapter sequences, and then aligned to the reference gene for extraction of mappable reads. VAULT extracts UMI sequence, followed by counting of the occurrence of each UMI, which reflects the number of reads in each UMI group. If a structured UMI (NNNNTGNNNN) is used in the experiment, the program will also check the UMI structure and separate them to perfect UMIs and wrong UMIs. Next, based on a user-defined threshold of minimum reads per UMI group, the program bins reads for eligible UMIs. The grouped reads will be subjected to alignment, followed by SNP and SV calling. After finishing all variant calling, a final data cleanup is performed to combine individual variant call files (VCF) together and filter the VCF. The number of reads in UMI groups and the corresponding UMI sequence will be written in the ID field of the VCF. Individual folders named after the UMI sequence will be saved to contain the alignment summaries and BAM files of every UMI group. VAULT supports both long-read data and single-end/ paired-end short-read data. The data analysis pipeline employs parallel computing for each UMI group, which avoids crosstalk during data analysis and accelerates the process. A typical analysis of 2.5 million long reads will take around four hours on a 32-core workstation.

**Fig. S1.**
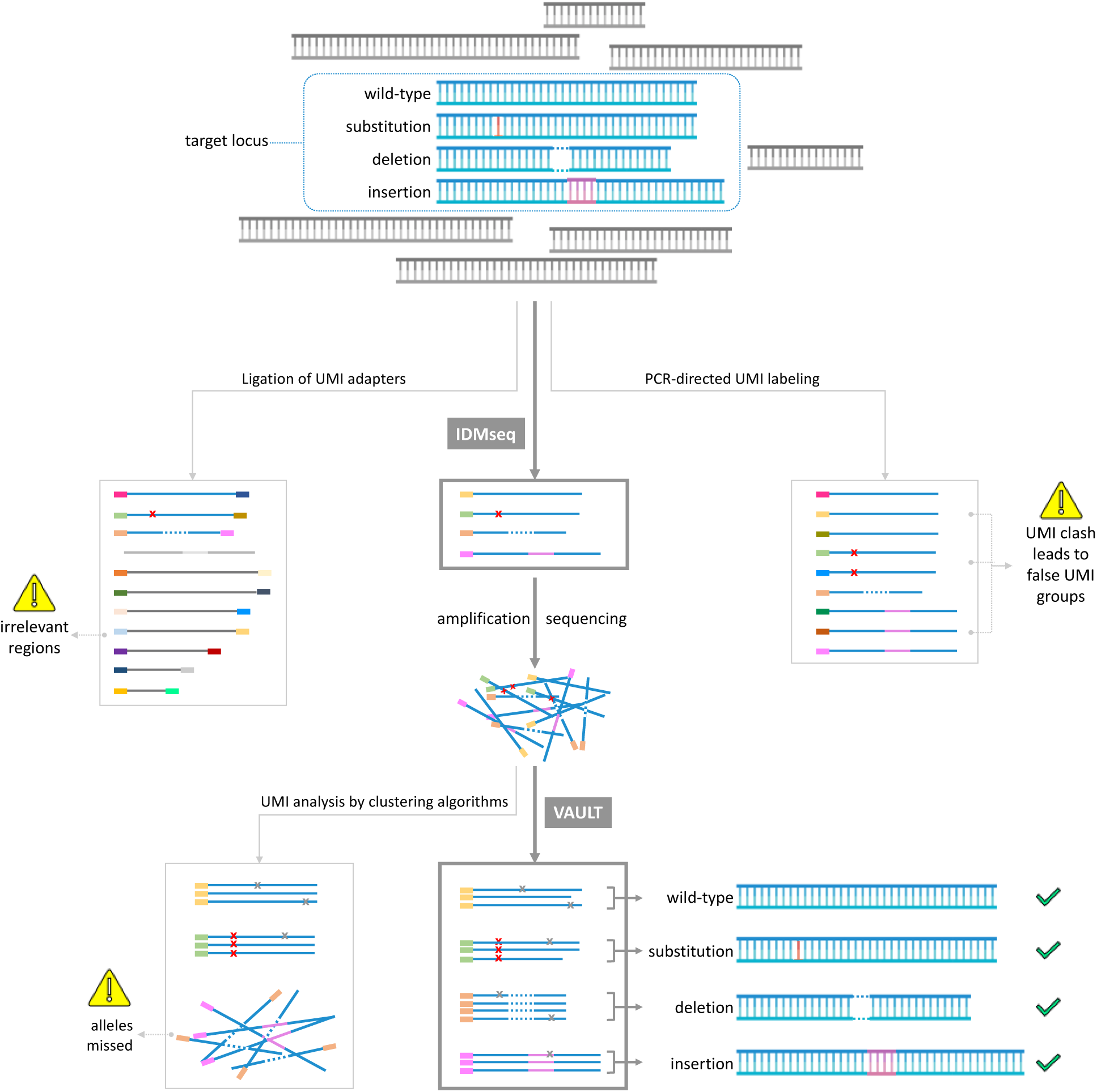
Advantages of IDMseq and VAULT as compared to current methods. A given population of cells (symbolized by the blue shape) may contain different alleles of a target locus, which accounts for a small proportion of the pool of genomic DNA. The first step of targeted molecular consensus sequencing is labeling of the variant alleles with UMI. Ligation-based and PCR-directed UMI labeling are two of the most widely used methods. Ligation-based UMI labeling will label irrelevant regions and the low efficiency of ligation will also omit a proportion of target alleles (greyed out in the middle left panel). PCR-directed UMI labeling is highly efficient but will result in UMI clashes (one original molecule labeled with multiple UMIs, leading to false UMI groups, middle right panel). IDMseq is the only method with high labeling efficiency and can faithfully retain the allele information (variants and frequency). After UMI labeling, the DNA with UMIs are amplified for sequencing in appropriated platforms (Illumina, Nanopore or PacBio). In the data analysis step, the algorithm needs to identify reads with the same UMI and use these to get the consensus sequence of the allele. This step is currently done with read-clustering algorithms that work well for fixed-length reads of short-read sequencing (e.g. Illumina). However, this strategy could miss reads with complex changes such as those uncovered by long-read sequencing, which prevents detection of deletions, insertions and complex structural variants (lower left panel). VAULT performs a BLAST-like strategy to locate UMI sequence in reads regardless of length and structure. VAULT analysis thus preserves the sequence information of all types of alleles and their frequency (lower middle and right).

**Fig. S2.**
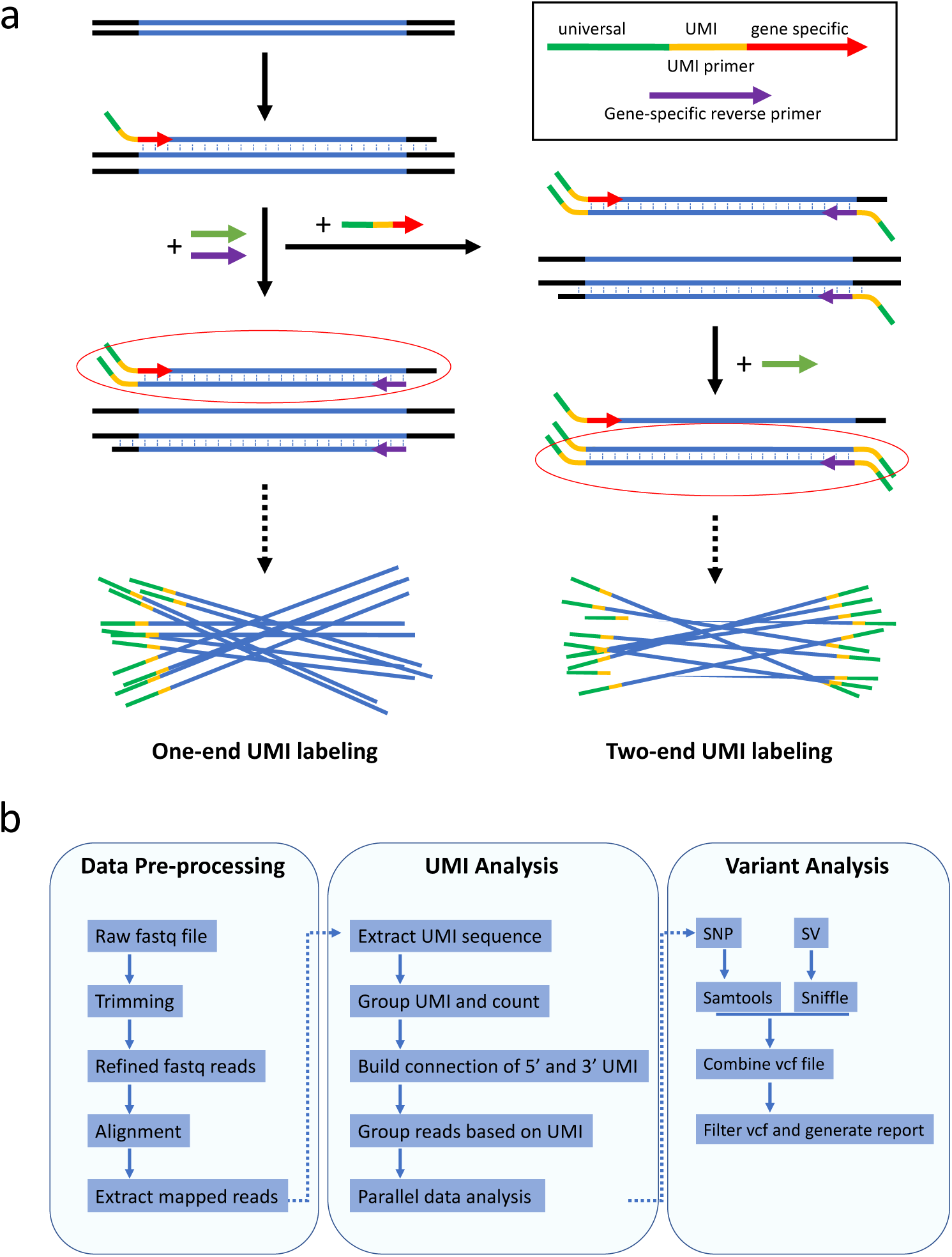
Overview of IDMseq and VAULT. **(a)** Schematic representation of UMI labeling. UMI primers are used to label individual DNA molecules with unique UMIs (one molecule is labeled with one UMI). It contains a 3’ gene-specific sequence, a UMI sequence, and a 5’ universal primer sequence. The 3’ gene-specific sequence is selected for its high specificity to the target gene. The middle UMI sequence consists of multiple random bases (denoted by Ns). The 5’ universal primer sequence is used to uniformly amplify all UMI-tagged DNA molecules. IDMseq is different from other UMI-based methods in that barcoding is achieved by a single round of primer extension rather than multiple cycles of PCR as commonly practiced_1,2_. For two-ended labeling, an additional round of primer extension with reverse UMI primers will be done after removing forward UMI primers. The UMI-labeled DNA will be further amplified by universal primers before sequencing. **(b)** Pipeline of VAULT analysis. During data per-processing, raw reads were filtered and mappable reads were extracted. After that, VAULT applies a BLAST-like strategy to locate UMI sequence in reads by searching for the known sequences of the universal primer and gene-specific forward primer. After that, VAULT bins reads according to UMI. The last steps of VAULT are variant calling for both SNVs and large SVs and report generation.

**Fig. S3.**
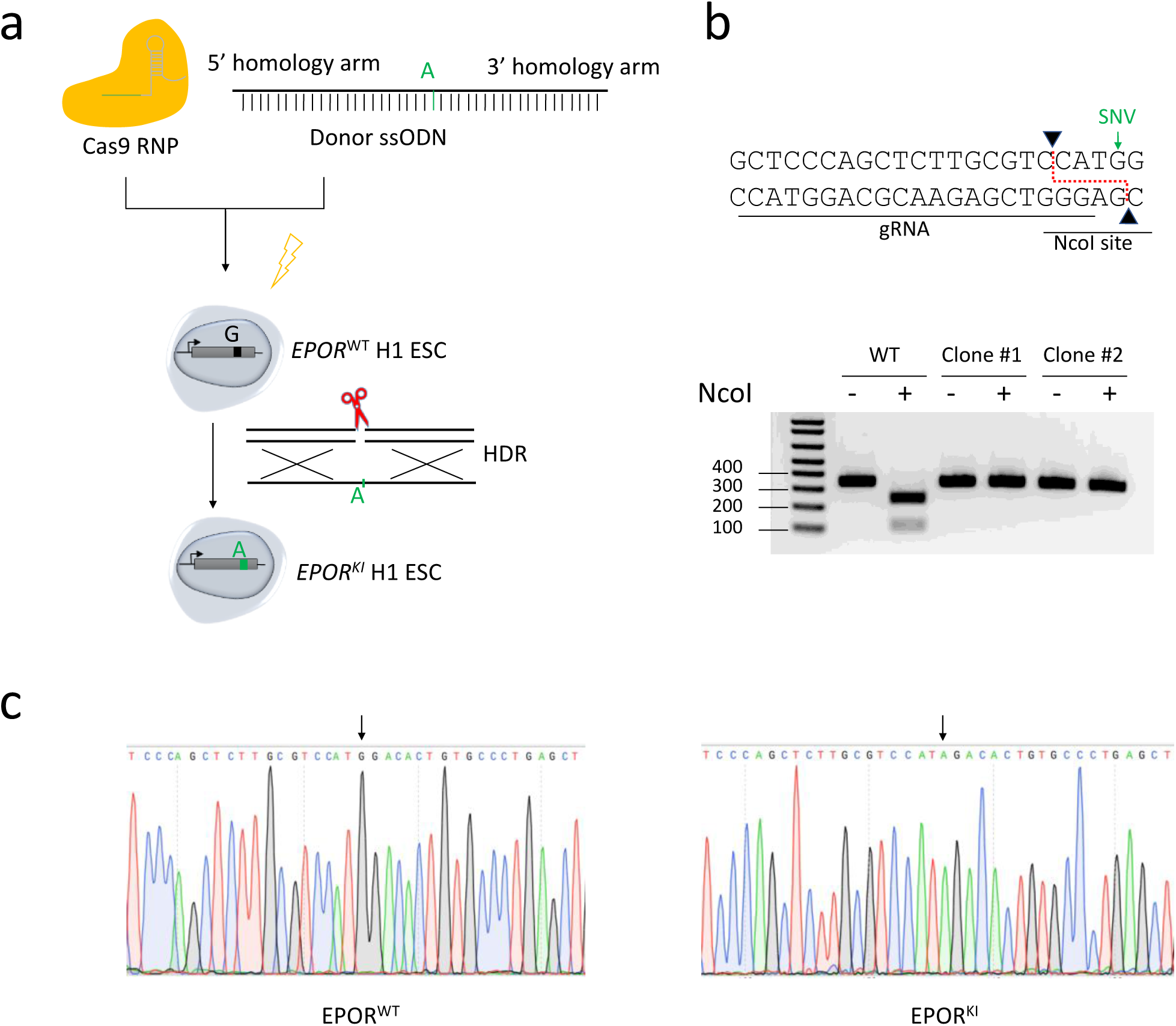
Generation of isogenic knock-in hESCs using CRISPR-Cas9. **(a)** Schematic representation of the experimental design. Cas9 RNP and ssODN were electroporated to H1 ESCs to generate homozygous G>A single-base substitution in the EPOR gene. **(b)** Schematic of the Cas9 target site and the NcoI restriction site. The gel image shows restriction enzyme digestion assay used to identify the knock-in hESC clones. Wild-type *EPOR* gene contains a NcoI site and thereby can be digested. The Knock-in allele will lose the NcoI site and cannot be digested. **(c)** Sanger sequencing results confirming the knock-in SNV. The homozygous SNV is indicated by the black arrow.

**Fig. S4.**
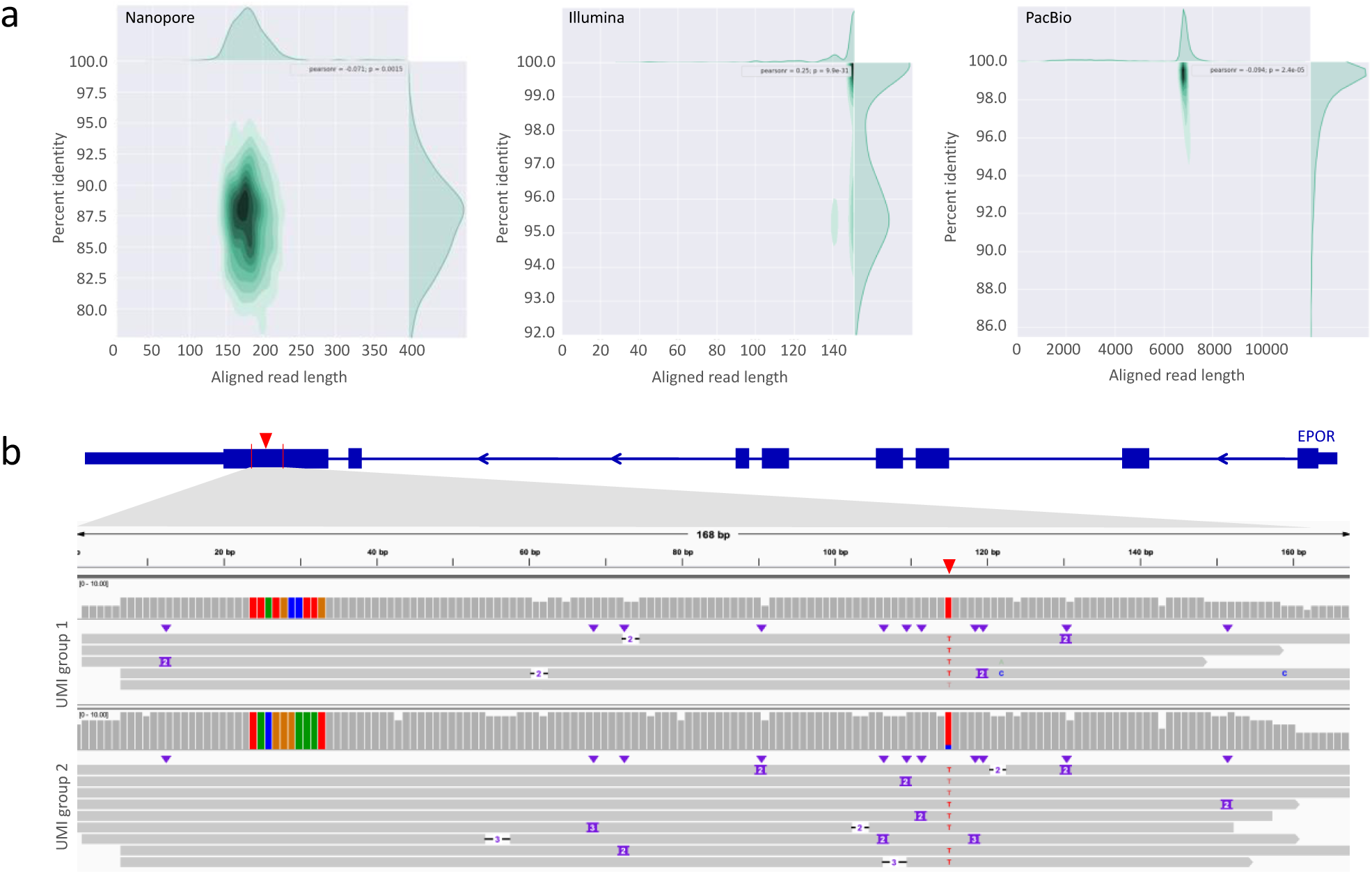
IDMseq for detection of the spike-in SNV. **(a)** Aligned read length vs. percent identity plot using kernel density estimation for Nanopore sequencing of the 1:10,000 population, Illumina sequencing of the 1:10,000 population, PacBio sequencing of the 1:1,000 population. **(b)** Two examples of Integrative Genomics Viewer (IGV) tracks of UMI groups in which the spike-in SNV in the 1:100 population was identified by Nanopore sequencing in conjunction with IDMseq and VAULT. The knock-in SNV is indicated by the red triangle in the diagram of the EPOR gene on top, and also shown as red “T” base in the alignment map. The gray bars show read coverage. The ten colored bars on the left side of the coverage plot represent the UMI sequence for the UMI group. Individual Nanopore reads within the group are shown under the coverage plot.

**Fig. S5.**
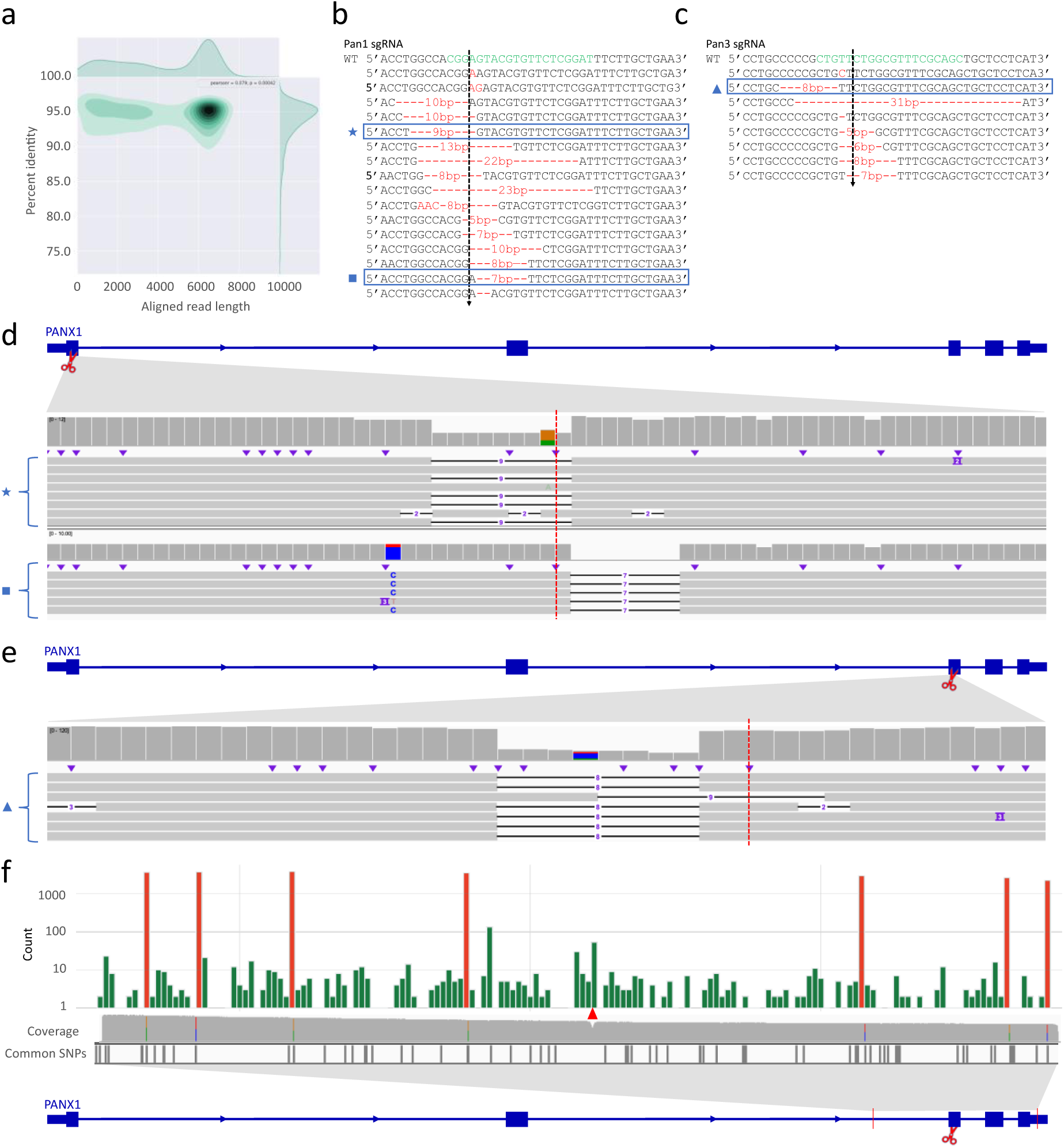
Analysis of CRISPR-Cas9 on-target mutagenesis. **(a)** Aligned read length vs. percent identity plot using kernel density estimation of Nanopore sequencing data of a 6595 bp region encompassing the Cas9 cleavage. **(b)** Individual alleles from Sanger sequencing of single-cell derived hESC clones after Cas9-directed mutagenesis in exon 1 of *PANX1*. Green letters indicate the gRNA sequence and the cleavage site is indicated by a dotted line. Red texts indicate insertion or deletion events. **(d)** IGV tracks of the UMI groups that shared the same deletion as alleles detected by Sanger sequencing of hESC clones (highlighted by a blue box in **(b)** and matching symbols). The red dotted line shows where Cas9 cuts. **(c)** similar to **(b)** but with the Pan3 sgRNA. **(e)** similar to **(d)** but with the Pan3 sgRNA. **(f)** Distribution of SNVs detected by IDMseq and VAULT in Pan3 edited hESCs. Somatic SNVs are shown in green, while the cell-line specific SNVs are shown in red. Somatic SNVs cannot be detected if variant calling is done en masse without UMI analysis (see the coverage track). Cell-line specific SNVs are detected in ensemble analysis (see colored lines in the coverage track) and most of them have been reported as common SNPs in dbSNP-141 database (Common SNPs track). The Cas9 cut site is indicated by the red triangle.

**Fig. S6.**
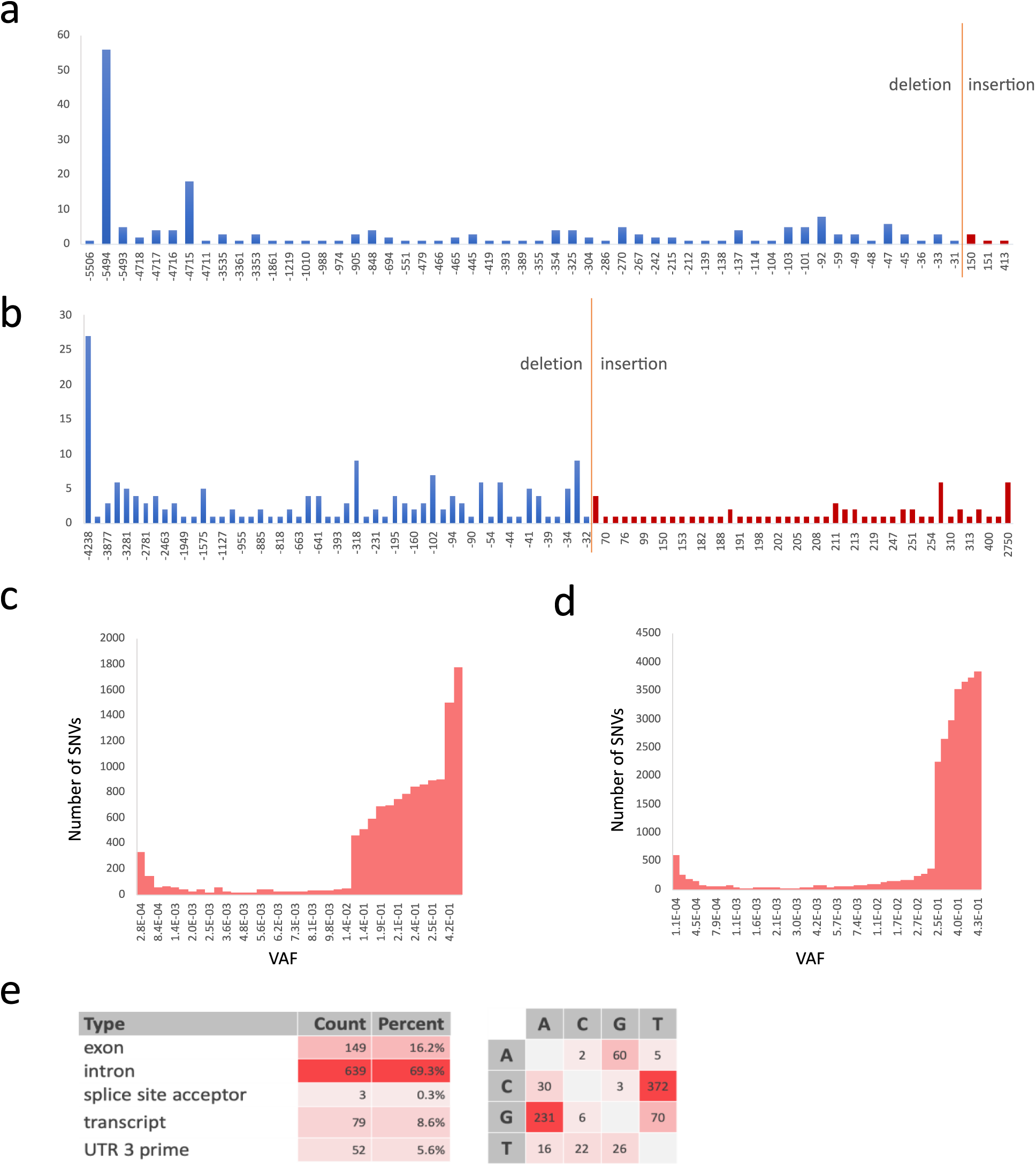
Analysis of variants in *PANX1* edited hESCs. **(a)** The frequency of deletions or insertions of different size detected in Pan1-edited hESCs. Certain deletions and insertions occur at disproportionally high frequencies. For example, a 5494 bp deletion was found in 56 UMI groups, which indicates a possible hotspot of Cas9-induced large deletion. **(b)** The frequency of different size deletions or insertions detected in Pan3-edited hESCs. Certain deletions and insertions occur at disproportionally high frequencies. For example, a 4238 bp deletion was found in 27 UMI groups, which indicates a possible hotspot of Cas9-induced large deletion. **(c, d)** Frequency distribution of the variant allele fraction of SNVs detected by IDMseq in Nanopore sequencing of the *PANX1* locus in Pan1-edited hESCs **(c)**, and Nanopore sequencing of the *PANX1* locus in Pan3-edited hESCs **(d)**. **(e)** Analysis of somatic mutations detected in Pan3-edited hESCs based on functional annotation and base change. The majority of base changes are G to A and C to T.

